# Comparative genomics and metagenomics analyses of endangered Père David’s deer (*Elaphurus davidianus*) provide insights into population recovery

**DOI:** 10.1101/073528

**Authors:** Xuejing Zhang, Cao Deng, Jingjing Ding, Yi Ren, Xiang Zhao, Shishang Qin, Shilin Zhu, Zhiwen Wang, Xiaoqiang Chai, Huasheng Huang, Yuhua Ding, Guoqing Lu, Lifeng Zhu

**Affiliations:** Nanjing Normal University, College of life Sciences, Nanjing 210046, China.; PubBio-Tech Services Corporation, Wuhan 430070, China.; Jiangsu Academy of Forestry, Nanjing, China.; Shanghai Majorbio Bio-pharm Biotechnology Co. Ltd., Shanghai, China.; DNA Stories Bioinformatics Center, Chengdu, 610021, China.; Jiangsu Dafeng Milu National Nature Reserve, Dafeng, China.; University of Nebraska at Omaha, Omaha, USA.

**Keywords:** Père David’s deer, comparative genomics, metagenomes, selective sweeping, population recovery

## Abstract

The milu (Père David’s deer, *Elaphurus davidianus*) has become a classic example of how highly endangered animal species can be rescued. However, the mechanisms that underpinned this population recovery remain largely unknown. As part of this study, we sequenced and analyzed whole genomes from multiple captive individuals. Following this analysis, we observed that the milu experienced a prolonged population decline over the last 200,000 years, which led to an elongated history of inbreeding. This protracted inbreeding history facilitated the purging of deleterious recessive alleles, thereby ameliorating associated threats to population viability. Because of this phenomenon, milu are now believed to be less susceptible to future inbreeding depression occurrences. SNP distribution patterns confirmed inbreeding history and also indicated sign of increased and increasing diversity in the recovered milu population. A selective sweep analysis identified two outlier genes (*CTSR2* and *GSG1*) that were related to male fertility. Furthermore, we observed strong signatures of selection pertaining to the host immune system, including six genes (*SERPINE1, PDIA3, CD302, IGLL1, VPREB3*, and *CD53 antigen*), which are likely to strengthen resistance to pathogens. We also identified several adaptive features including the over-representation of gene families encoding for olfactory receptor activity, a high selection pressure pertaining to DNA repair and host immunity, and tolerance to high-salt swamp diets. Moreover, glycan biosynthesis, lipid metabolism, and cofactor and vitamin metabolism were all significantly enriched in the gut microbiomes of milu. We speculate that these characteristics play an important role in milu energy metabolism, immunity, development, and health. In conclusion, our findings provide a unique insight into animal population recovery strategies.

## Introduction

Milu were once widely distributed in the swamps of East Asia, and they were predominantly found in China (**Figure 1AB**, **Supplementary Fig. S1**). This species was first introduced to west in 1866 by Armand David (Père David)(Cao 2005), and subsequently became extinct in its native China in the early 20^th^ century(Cao 2005). Fortunately, between 1894 and 1901, Herbrand Arthur Russell (the 11^th^ Duke of Bedford), acquired the few remaining deer (18 individuals) from European zoos. These individuals were nurtured at Woburn Abbey in England(Cao 2005) (**Figure 1C**) and the current world population was derived from this herd(Cao 2005). In the mid-1980s, 77 individuals were reintroduced to captive facilities in China(Cao 2005; Jiang and Harris 2008), and populations were established in Beijing, Dafeng, Tianezhou and Yuanyang (**Figure 1C**). Since then, the populations have rapidly expanded, and the milu have managed to overcome the genetic bottleneck of inbreeding. The repopulation of milu is now deemed a classic example of how a highly endangered species can be rescued. However, the mechanisms that underpin this population recovery remain largely unknown.

**Figure 1.**
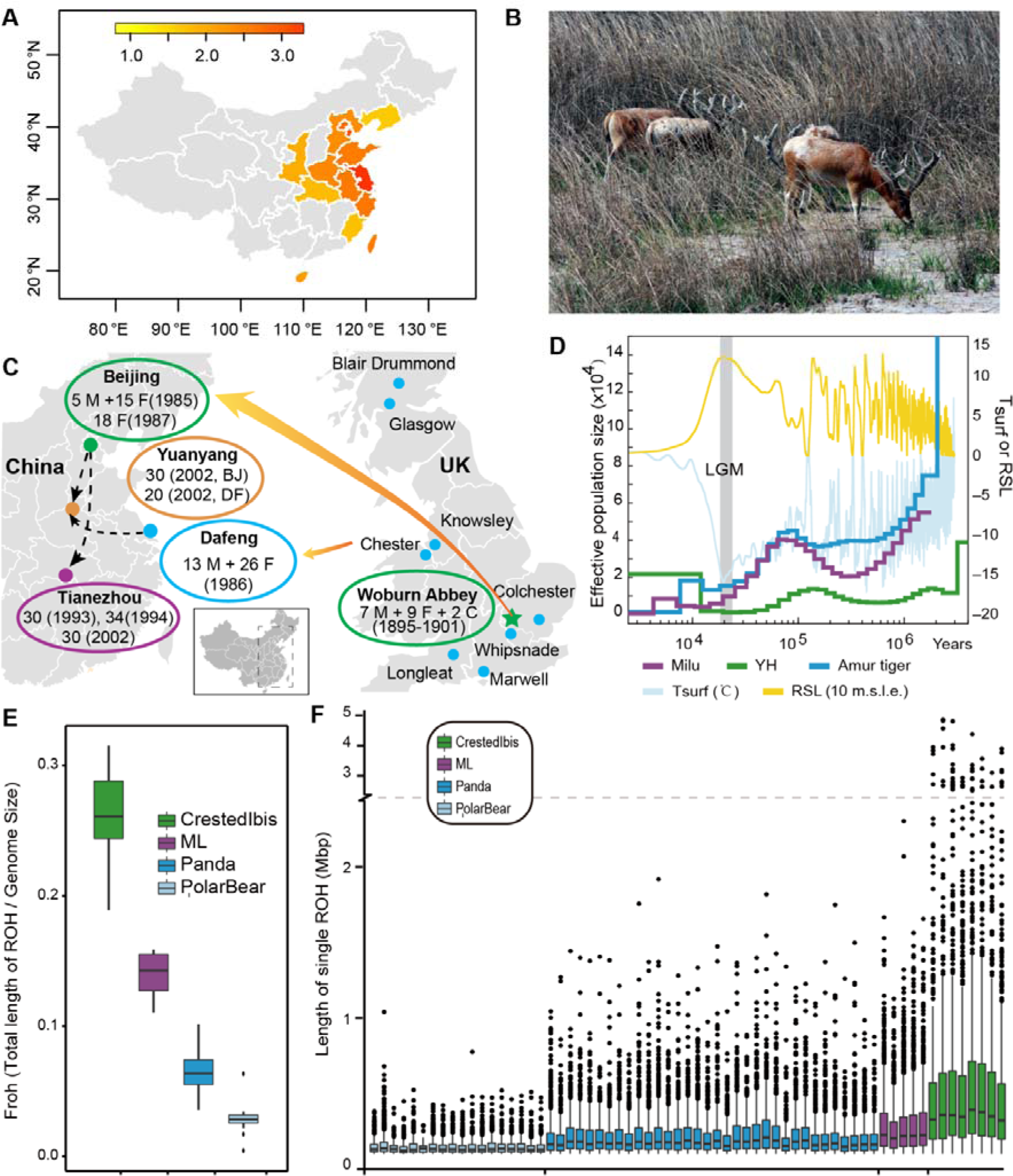
History of milu. **A**, Palaeogeographic distribution history of wild milu in China. The data for milu fossils were adopted from Cao^1^. The color relates to the density of the fossils in specific provinces, and the density was calculated as the number of fossils per million square kilometers. **B**, Forage selection in coastal shoal habitat of milu in Dafeng Milu Natural Reserve, Jiangsu, China. **C**, Large-scale reintroduction programs since 1985. C, fawn; F, females; M, males. **D**, Demographic history of the milu. The history of the milu population and climate change spans from 3 KYA to 4 MYA. We used the default mutation rate of 1.5×10^−8^ for baiji (μ) and an estimation of 6 years per generation (g). The last glacial maximum (LGM) is highlighted in grey. Tsurf, atmospheric surface air temperature; RSL, relative sea level; 10 m.s.l.e., 10 m sea level equivalent. **E**, Box plot of *Froh* for milu, crested ibis, panda, and polar bear populations. *Fron* denotes the proportion of total ROH length. **F**. Box plot of length of ROH in each individual from milu, crested ibis, panda, and polar bear.

### Results and Discussion

We sequenced and analysed the milu genome and performed whole-genome re-sequencing for five another individuals. The assembled genome (2.58 GB; ∼114-fold coverage) had a scaffold N50 value of 2.85 Mb (**Supplementary Table S2**). Assembly quality assessment was performed by aligning the transcripts from *Odocoileus virginianus* (white-tailed deer, WTD) and *Cervus nippon* (Chinese Sika deer, CSD) to the scaffolds of milu (>93.9% and >97.6% coverage, respectively) (**Supplementary Table S3**) and a core eukaryotic gene set (>92.0% conserved genes). We observed that repetitive sequences occupied 39.84% of the whole assembly (**Supplementary Table S4-S5**), and 22,126 protein-coding genes were predicted by combining *de novo* and evidence-based gene predictions (**Supplementary Table S6**).

Milu had been raised in enclosures for more than 1,200 years, with supplementation occurring through the introduction of wild individuals(Li et al. 2011). This resulted in a prolonged genetic bottleneck with low resultant genetic diversity. Results generated using the Pairwise Sequentially Markovian Coalescent (PSMC) model(Li and Durbin 2011) validated this hypothesis (**Figure 1D**, **Supplementary Fig. S33**). After the Last Glacial Maximum (LGM, ∼20 thousand years ago/KYA)(Yokoyama et al. 2000), it is likely that milu suffered from the effects of climate change, over-hunting and/or habitat loss. Indeed, milu populations diminished, and there was a tendency towards continuous decreases. This is further evidenced by fossil records and associated literary records(Cao 2005).

Reduced population sizes increase the opportunity for inbreeding. The protracted existence of small populations along with more recent declines resulted in high levels of milu inbreeding. When related individuals mate, the offspring carry long stretches of homozygous genome. Thus, the detection of runs of homozygosity (ROH) is a practical approach for estimating inbreeding at the individual level(Kim et al. 2013; Zhou et al. 2014) (**Supplementary Table S36**). When compared with 34 giant panda genomes(Zhao et al. 2013), 18 polar bear genomes(Liu et al. 2014) and eight Crested ibis(Li et al. 2014) genomes, we observed that the *Froh* (ROH length / Genome effective length) of milu ranged from 0.11 to 0.16. These values are much higher than those exhibited by the well-known panda (from 0.04 to 0.10) and polar bear (from 0.004 to 0.064), which are less prone to occurrences of inbreeding. However, the milu *Froh* values are lower than those exhibited by the previously critically-endangered crested ibis (from 0.19 to 0.32), which experienced a more recent and severe genetic bottleneck(Li et al. 2014) (**Figure 1E**). Length distribution of ROH also provides information about the timing of major inbreeding events. Long ROH are most likely derived from a recent ancestor; shorter ones, from a more distant ancestor(Curik et al. 2014). As revealed in **Figure 1F**, the milu has a medium average ROH length when compared with the crested ibis, the panda and the polar bear. The crested ibis contains an elongated ROH (longer than 1M), which is consistent with the fact that current crested ibis populations are derived from seven individuals approximately 40 years ago(Li et al. 2014). The milu harbors an increased average ROH length compared with the pandas and polar bears; however, this value is shorter than those observed for crested ibis. This would suggest that the time of major milu inbreeding event occurred prior to that of crested ibis but after those of panda and polar bear. These data confirm the existence of a prolonged reduced milu population.

Another major threat to small and endangered populations involves the loss of genetic diversity(Frankham 2005; Steiner et al. 2013). Small populations are susceptible to genetic drift and fixation, and these phenomena can be accelerated by inbreeding(Saccheri et al. 1998; Keller and Waller 2002; Steiner et al. 2013). We observed that genetic diversity was lower in the milu than in the panda, with a heterozygosity rate of 0.51 per kilobase pair in the milu, versus 1.32 per kilobase pair in the panda (**Supplementary Table S25**). Comparison with other endangered animals that experience, or have experienced, ongoing or recent population bottlenecks, indicated that this value was similar to that of mountain gorillas (Xue et al. 2015) (0.64×10^−3^) but slightly higher than that of the crested ibis (0.36×10^−3^, **Figure 2A**), Chinese alligator(Wan et al. 2013)(0.15×10^−3^) and baiji(Zhou et al. 2013) (0.12×10^−3^). In addition, patterns of SNP density distributions were explored by fitting a two-component mixture model to the observed SNP densities using the expectation-maximization algorithm(Hacquard et al. 2013) (**Figure 2B**, **Supplementary Table S30-S33**). Half of the milu genome harbored only less than 5% of the called SNPs, and the mean heterozygosity of these low SNP density regions was 0.03 per kilobase, a value that was similar to that observed in crested ibis but much lower than those observed in panda and polar bear, reflecting more recent inbreeding history in milu and crested ibis. However, the mean heterozygosity in the other half of the milu genome was 1.26 per kilobase, which was similar to that observed in panda but higher than that observed in crested ibis, indicating a stronger sign of increased diversity in the recovered milu population than crested ibis population. Generally, the occurrence of heterozygosity in exons is reduced due to selective constraints(Li et al. 2014). However, the ratio of exon heterozygosity to genome heterozygosity in the milu and crested ibis is higher than that observed for the panda and polar bear (**Figure 2C**, **Supplementary Table S25**). There are two possible explanations for this finding. First, it is possible that the milu and crested ibis experienced a slower rate of loss of genetic diversity in exons during inbreeding. Second, a rapid increase in the diversity of exons in recovered milu and crested ibis populations, following the occurrence of severe genetic bottlenecks, may have resulted in greater genetic diversity in these genetic regions. Inbreeding depression is a major force affecting the evolution and viability of small populations in captive breeding and restoration programs(Saccheri et al. 1998; Keller and Waller 2002; Steiner et al. 2013). Deleterious mutations tend to accumulate in associated populations due to reduced selective strength(Saccheri et al. 1998; Steiner et al. 2013). We observed that the milu exhibits a relatively low percentage of deleterious variants compared to other healthy or recovered populations (**Figure 2D**). This is consistent with a low effective population size (Ne) and the occurrence of inbreeding(Xue et al. 2015). In these populations, alleles occur more frequently in the homozygous state, and because deleterious variants are more likely to be pronounced, they are less likely to persist in the population (even if recessive)(Xue et al. 2015). Therefore, populations, such as the milu, that have experienced reduced population sizes for prolonged periods may be less susceptible to future inbreeding depressions because they have been purged of deleterious recessive alleles. Consequently, these populations are more likely to recover from future severe genetic bottlenecks.

**Figure 2.**
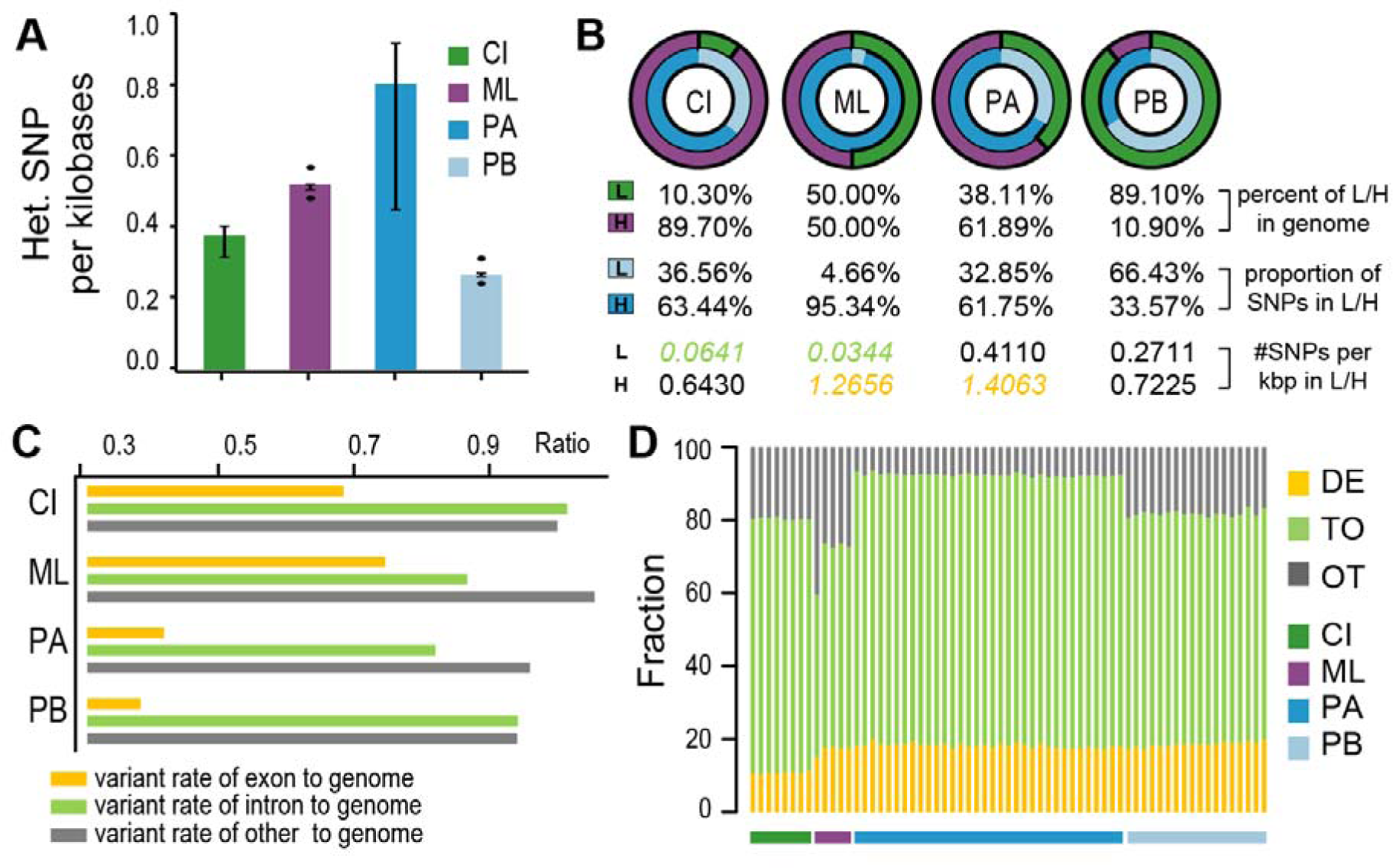
Genetic diversity of milu and other animals. **A**. Box plot of heterozygosity from milu, crested ibis, panda, and polar bear individuals. Only heterozygous SNPs were included. CI, Crested Ibis; ML, Milu; PA: Panda; PB: Polar bear. **B**. Bias distribution of SNPs in animal genomes. Each circle denotes one species as (**A)**. L, low SNP density region; H, high SNP density region; kbp, kilobase; the proportion of total length of L and H regions in whole genome are green and purple; the proportion of SNP number in L and H region to total SNP number in both L and H regions are light blue and blue. **C**. Ratio of heterozygosity in each genomic element. The genomes were subdivided into three regions – exons, introns and other (regions that were neither exons nor introns). Then, heterozygosity in each type of genomic element was compared to heterozygosity of whole genome. **D**. Classification of missense variants. DE: deleterious; TO: Tolerated; and OT: Other.

Because of the prolonged history of captivity, reduced population size, and inbreeding associated with the milu, the study of adaptive evolution following exposure to these conditions is imperative in our efforts to prevent further future bottlenecks. We investigated adaptive evolution in the milu by analyzing the composition of several protein domains, and the expansion and contraction of a number of gene families^14,19,20^. We also investigated lineage-specific accelerated evolving GO categories(Sequencing and Consortium 2005; Bakewell et al. 2007; Qiu et al. 2012) and PSGs(Qiu et al. 2012; Zhou et al. 2013; Yim et al. 2014) (**Supplementary Materials**). A functional analysis of the milu-specific expansion domains (**Supplementary Table S13**) showed that a large proportion of such domains is related to translation machinery. Notably, HSP90 genes in milu show a remarkable expansion in cytosolic members (*HSP90AA* and *HSP90AB*), especially the inducible HSP90AA1 and HSP90AA2 forms (**Supplementary Table S14, Supplementary Fig. S15**). The Hsp90 protein (PF00183) is important in stress response and has a capacity to buffer underlying genetic variation(Yeyati et al. 2007). Upon analysis of gene family numbers, we identified 835 and 4,584 gene families that expanded and contracted in the milu, respectively. In other mammals, it was observed that gene families expanded (p<0.01, **Figure 3A**). The more pronounced expanded families were significantly over-represented (**Supplementary Table S11**) by genetic elements pertaining to ‘olfactory receptor activity’ (P=3.29 × 10^65^), detection of chemical stimulus involved in sensory perception of smell (P=2.00 × 10^47^), ‘ATPase activity’ (P=6.26 × 10^7^), ‘platelet dense granule membranes’ (P=5.65 × 10^14^), chloride channel activity (P=7.92 × 10^6^), antigen processing and presentation of peptide antigen via MHC class I (P=3.54 × 10^3^), cellular response to interferon-gamma (P=1.35 × 10^3^), sperm mitochondrial sheath (P=9.85 × 10^3^). These functional groups might play important roles in milu’s behavior, development, immune and breeding. For example, much of the cellular response to interferon-gamma can be described in terms of a set of integrated molecular programs underlying well-defined physiological systems; and the induction of efficient antigen processing for MHC-mediated antigen presentation, which play clearly defined roles in pathogen resistance(Boehm et al. 1997). We also identified 26, 25, and 17 GO categories that demonstrated a significantly elevated pairwise number of non-synonymous substitution (*A*) values in the milu in the comparison of milu-cow-human, milu-TA-human and milu-baiji-human, respectively; while 14, 21, and 30 GO categories were elevated in cow, TA and baiji, respectively. In reference to the milu, the accelerated evolving GO categories were predominantly found to be involved in DNA repair, gene expression, protein modification, development, immunity, excretion, and responses to insulin stimuli (**Figure 3B**, **Supplementary Table S17-S18, Supplementary Fig. S17-S18**). Furthermore, 455 PSGs were identified using the likelihood ratio test implemented in PAML(Yang 2007) (**Supplementary Table S19**). These PSGs were enriched for genes involved in DNA repair, RNA metabolic processes, cellular protein modification processes, nitrogen compound metabolic processes, TLR 3 signaling pathways, regulation of development processes, and regulation of cytokine production (**Supplementary Table S20, Supplementary Fig. S19**).

**Figure 3.**
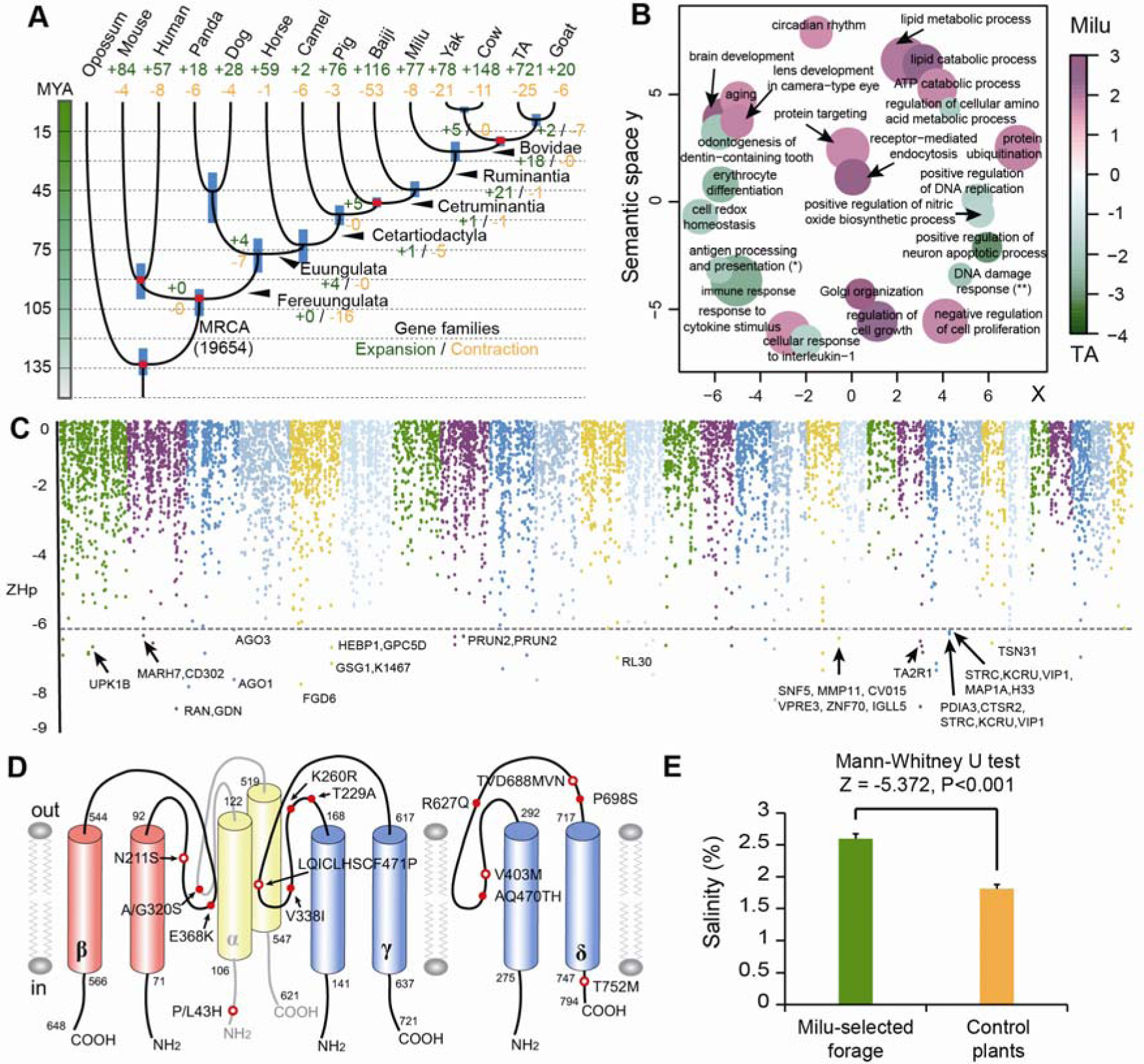
Adaptive evolution in the milu genome. **A**. Phylogenetic position of milu relative to other mammals. The branch lengths of the phylogenetic tree are scaled to demonstrate divergence time. Tree topology is supported by a posterior probability of 1.0 for all nodes. The blue bars on the nodes indicate the 95% credibility intervals of the estimated posterior distributions of the divergence times. The red circles indicate the fossil calibration times used for setting the upper and lower bounds of the estimates. The number of significantly expanded (green) and contracted (orange) gene families is designated on each branch. MRCA, most recent common ancestor. **B**. Lineage-specific accelerated evolving GO categories of biological process using the number of non-synonymous substitutions. **C**. Summary of selective sweep analysis. The negative end of the *ZHp* distribution presented along pseudo-chromosomes 1–29. The horizontal dashed lines indicate the threshold at *ZHp* = -6. Genes residing within 20 kb of a window with *ZHp* ≤ -6 are indicated by their gene names. **D**. Red dot, milu-specific SAPs (single amino acid polymorphisms); red circle, damaging milu-specific SAPs predicted by PPH2. **E**. The salinity of forage plants in Dafeng Milu Natural Reserve.

In small captive populations, genetic adaptation to artificial environments can also occur, through processes including selective sweeps(Rubin et al. 2010; Rubin et al. 2012). We searched the genome for regions with high degrees of fixation, and the distributions of observed *Hp* values and the Z transformations of *Hp*, *ZHp*, are shown in **Figure 3C**. In the genome-wide screen, 30 distinct gene loci showed a *ZHp* value lower than −6. Among the outliers derived following this analysis, we observed two genes that are related to male fertility, *CTSR2* (a.k.a. *CATSPER2*, cation channel sperm-associated protein 2) and *GSG1*(Germ cell-specific gene 1 protein). *CTSR2* complexes with other family members to form a calcium permeant ion channel, which plays a primary role in the regulation of sperm motility(Quill et al. 2003). *GSG1* colocalized with testis-specific poly(A) polymerase (*TRAP*) during spermiogenesis, and the interaction between *TPAP* and *GSG1* may be related to morphological alterations that occur during spermiogenesis (the transformation of round spermatids to elongating spermatids)(Choi et al. 2008). This may imply that potential selection of breeding stocks occurred in the milu population, thereby supporting the prolonged captive history of the latter. Interestingly, the gene family of sperm mitochondrial sheath (P=9.85 × 10^−3^) was significant expanded in Milu genome. The mature sperm tail has several accessory structures, including a mitochondrial sheath, outer dense fibers and a fibrous sheath, and (Holstein 1976). Studies with gene knockout mice have proven that precisely regulated mitochondrial sheath formation is critical for sperm motility and fertility(Bouchard et al. 2000; Miki et al. 2004).

We also observed strong signatures of selection in relation to host immunity, including six genes (*SERPINE1, PDIA3, CD302, IGLL1, VPREB3*, and *CD53 antigen*), which may strengthen host resistance to pathogenic infection. Another interesting signature of positive selection was the *TAS2R* locus (**Figure 3C**). The *TAS2R* locus controls bitter taste sensitivity, including sensitivity to saccharin, quinine, and salicin(Deshpande et al. 2010). Moreover, we also found the significant gene family expansion on chloride channel activity in Milu genome, which mediates salt and liquid movement(Sheppard and Welsh 1999). By scanning milu-specific single amino acid polymorphisms (SAPs) in salt-sensitive ENaCs (epithelial sodium channels)(Chandrashekar et al. 2010), we identified 14 SAPs associated with *SCNN1A*, *SCNN1B*, *SCNN1G*, and *SCNN1D*(**Supplementary Table S39, Supplementary Fig. S34-S37**). Eight SAPs were predicted to influence channel function, thereby affecting salt-sensation and sodium absorption (**Figure 3D**). Historically, milu were widely distributed in the eastern coastal regions of China(Cao 2005) (**Figure 1A**). Currently, the largest captive and wild release populations live in Dafeng Natural Reserve, in the eastern coastal shoal region of China (**Figure 1C**). The salinity of the main diet of these individuals is significantly higher than for inland populations (**Figure 3E**, **Supplementary Table S40-S41**). Thus, the occurrence of polymorphisms in loci that are related to bitter and salt tasting sensations may explain the adaption of the milu to high-salt diets in swamp.

Symbiotic gut microbes play important roles in host nutrition, development, immunity, and health in animals(Ley et al. 2008). Metagenomic analysis of 10 milu gut microbial genomes and 39 mammalian microbial genomes (including whale, dolphin, carnivore, omnivore and herbivore genomes)(Muegge et al. 2011; Sanders et al. 2015) was performed using the MG-RAST online server(Meyer et al. 2008) (**Supplementary Table S42**). This analysis revealed functional enrichment of sodium transportation in milu gut microbes. Factors that were affected by this phenomenon included the Sodium transport system ATP-binding protein, Adenosinetriphosphatase, and Transcriptional regulatory protein NatR (**Figure 4A-C**). These occurrences may reflect an adaptation to a high salinity diet. Moreover, glycan biosynthesis, lipid metabolism, cofactor and vitamin metabolism (including folate biosynthesis, thiamine biosynthesis and vitamin B6 metabolism), and biosynthesis of other secondary metabolites (including penicillin and cephalosporins) were also significantly enriched in milu gut microbes (**Figure 4D-G**). It is possible that these reactions participate in host immunity, development, and health.

**Figure 4.**
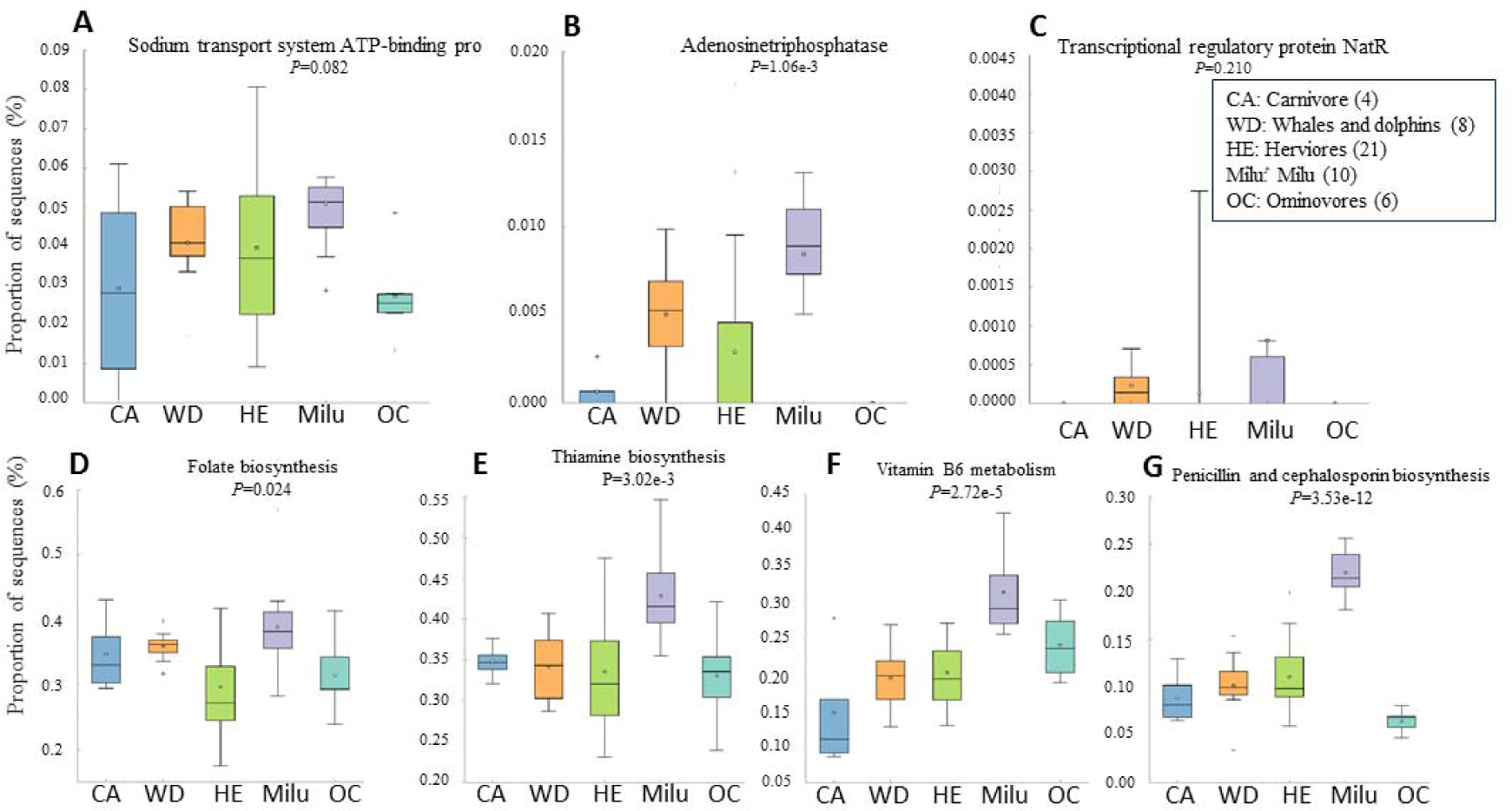
The comparative metagenomic analysis of 10 milu gut microbial genomes and another 39 mammalian genomes (including genomes from whales, dolphins, carnivores, omnivores and herbivores). **A-C**, the genes coding for putative enzymes related to the sodium transport system, including Sodium transport system ATP-binding protein, Adenosinetriphosphatase, and Transcriptional regulatory protein, NatR. **D-G**, the genes coding for putative metabolism of cofactors and vitamins (folate biosynthesis, thiamine biosynthesis and vitamin B6 metabolism), and biosynthesis of other secondary metabolites (including penicillin and cephalosporin biosynthesis). CA, carnivores. WD, whales and dolphins. HE, herbivores. OC, omnivores. The number in brackets represents sample size.

## Acknowledgments

This work was supported by grants from the National Natural Science Fund for outstanding young fund (31222009), National Natural Science Fund (31570489) and the Priority Academic Program Development of Jiangsu Higher Education Institutions (PAPD).

## Author Contributions

L.Z. conceived the study, L.Z. headed and Y.R managed the sequencing project, X.Z., and J.D prepared sequencing data, L.Z., C.D. and Z.W. coordinated the bioinformatics activities, L.Z., C.D., X.Z., S.Z., Z.W., S.Q. and X.C. designed experiments and analyzed the data, S.H., G. L., and Y.D. participated in project design, L.Z., C.D. and G. L. wrote and edited the manuscript with input from all other authors. All authors have read and have approved the manuscript.

## Author Information

The E. davidianus whole-genome sequences are deposited in GenBank under accession number JRFZ00000000.The 10 metagenomes of Milu gut microbes were submitted to MG-Rast, and the accession number were 4693474.3, 4693473.3, 4693472.3, 4693453.3, 4693450.3, 4693448.3, 4693446.3, 4693445.3, 4693207.3 and 4693196.3.Reprints and permissions information is available at www.nature.com/reprints. The authors declare no competing financial interests. Readers are welcome to comment on the online version of the paper. Correspondence and requests for materials should be addressed to L.Z.

## Online Methods

### Genome sequencing and assembly

DNA from blood samples acquired from an adult female milu in Dafeng Milu Natural Reserve was used for *de novo* sequencing. Samples from an additional five animals were utilized for resequencing. Libraries with different insert sizes were constructed at Majorbio (Shanghai), and the insert sizes of the libraries were 180 bp, 500 bp, 800 bp, 3 kb, 5 kb, 8 kb, and 10 kb. The libraries were sequenced using a HiSeq2000 instrument. The other five resequencing samples were sequenced with read and insert lengths of 101 bp and 500 bp, respectively.

Whole-genome shotgun assembly of the milu was performed using the short oligonucleotide analysis 316 package, SOAP *denovo*(Li et al. 2010). After filtering the reads, short-insert size library data were used to construct a *de Bruijn* graph without paired-end information. Contigs were constructed by merging the bubbles and resolving the small repeats. All qualified reads were realigned to contig sequences and paired-end relationships between the reads of allowed linkages between the contigs. We subsequently used the relationships, step by step, from the short-insert size-paired ends and the long-distance paired-ends to construct scaffolds. Gaps were then closed using the paired-end information to retrieve read pairs in which one end mapped to a unique contig and the other was located in the gap region. Assembly quality was assessed by aligning the assembled WTD(Malenfant et al. 2014) and CSD(Yao et al. 2012a; Yao et al. 2012b) transcripts with the milu scaffolds and by using a core eukaryotic gene mapping method(Parra et al. 2007).

### Genome annotation

Transposable elements in the milu genome were identified by a combination of homology-based and *de novo* approaches. Tandem repeats were identified using Tandem Repeat Finder(Benson 1999). Interspersed repeats were characterized by homolog-based identification using RepeatMasker open-4.0.3(Smit et al. 1996) and the repeat database, Repbase^2^. Repeated proteins were identified using RepeatProteinMask and the transposable elements protein database. *De novo* identified interspersed repeats were annotated using RepeatModeler(Price et al. 2005), and LTR_FINDER(Xu and Wang 2007) was used to identify the LTRs; these results were used to generate the *de novo* repeat libraries, and then RepeatMasker was run once more against the *de novo* libraries. All repeats identified in this manner were included in the total count of interspersed repeats.

The milu protein-coding genes were annotated following the use of a combination of homolog gene prediction and *de novo* gene prediction tools. For homolog gene prediction, the protein sequences from cow, yak, goat, TA, and human were mapped to the genome using tBLASTn(Altschul et al. 1990), and GeneWise(Birney et al. 2004) was used to predict the gene model based on the alignment results. *De novo* gene prediction was performed using GENSCAN(Burge and Karlin 1997), AUGUSTUS(Stanke et al. 2006), and GLIMMERHMM(Majoros et al. 2004) based on the repeat-masked genome. Then, EVM(Haas et al. 2008) and MAKER(Cantarel et al. 2008) were applied to integrate the predicted genes. Finally, manual integration was performed to construct the final gene set. We searched the final gene set against the KEGG(Kanehisa and Goto 2000), SwissProt(Bairoch and Apweiler 2000), and TrEMBL(Bairoch and Apweiler 2000) protein databases to identify gene functions. The gene motifs and domains were determined using InterProScan(Zdobnov and Apweiler 2001) following analysis of public protein databases, including ProDom, PRINTS, PFAM, SMART, PANTHER and PROSITE. All genes were aligned against the KEGG pathway database(Kanehisa and Goto 2000), and the best match for each gene was identified. The GO IDs for each gene were obtained from the corresponding InterPro entries. We also mapped milu proteins to the NCBI nr database and retrieved GO IDs using BLAST2GO(Conesa et al. 2005).

### Genome evolution

Orthologous groups were constructed by ORTHOMCL v2.0.9. Phylogenetic tree inference and divergence time estimation was conducted based on fourfold-degenerate sites of single-copy gene families. Significantly expanded and contracted gene families were identified by CAFE(De Bie et al. 2006). Molecular evolution analyses were performed using the framework provided by the PAML4.7 package. Please see Supplementary information for more detailed methodologies.

### Detection of variants

For the individual that was used for *de novo* sequencing, we used the BWA(Li and Durbin 2009) program to remap the pair-end (180 bp, 500 bp, and 800 bp) clean reads to the assembled scaffolds. After merging the BWA results and sorting alignments (using the leftmost coordinates) and removing potential PCR duplicates, we used SAMtools(Li et al. 2009) mpileup to call SNPs and short InDels. We applied vcfutils.pl varFilter (in SAMtools) as the filtering tool with parameters ‘*-Q 20 -d 6 -D 86*’. Then, homologous SNP positions were extracted and further filtered, to disqualify SNPs that may have resulted from errors due to assembly and/or mapping. The heterozygosity rate was estimated as the density of heterozygous SNPs for the whole genome, gene intervals, introns, and exons, respectively. For the five resequencing milu individuals, variants were identified using similar methods, except that the filtering parameter used by vcfutils.pl varFilter was ‘*-Q 20 -d 6 -D 75*’.

Whole genome re-sequencing data from 34 giant panda genomes(Zhao et al. 2013), and eight crested ibis(Li et al. 2014) genomes were downloaded from the NCBI SRA database, and BAM files were generated using identical methods to those used for milu individuals. Next, the bam files for each species were processed using the mpileup module in samtools and the following parameters; ‘-q 1 -C 50 -g -t DP, SP, DP4 -I -d 250 -L 250 -m 2 -p’. The associated variants were called and filtered using the varFilter module of vcfutils.pl (parameters ‘*-Q 20 -d 10 -D 50000 –w 5 -W 10*’ for panda, and ‘-Q 20 -d 5 -D 4000 -w 5 -W 10’ for crested ibis). Finally, variants from each individual were generated by filtering positions with low depth (‘<3’ for panda, and ‘<5’ for crested ibis). The SNP positions in 18 polar bear genomes(Liu et al. 2014) were extracted from variant files downloaded from GigaDB(Sneddon et al. 2012). SNPs were annotated using snpEff (Cingolani et al. 2012). To estimate how the functional changes for proteins in milu/panda/polar bear/crested ibis differed from those in humans, we evaluated the likely effect of a mutation in humans relative to the milu/panda/polarbear/crested ibis alleles as either neutral or deleterious using SIFT(Ng and Henikoff 2003).

### Demographic history reconstruction and ROH identification

Demographic histories of the milu were reconstructed using the Pairwise Sequentially Markovian Coalescent (PSMC) model(Li and Durbin 2011). The mutation rate (μ^) was set to 1.5×10^ -8 and the generation time (g) was set to 6 years. We identified the ROH for each individual using the runs of homozygosity tool in PLINK (v.1.07)(Purcell et al. 2007) with adjusted parameters (--homozyg-window-kb 0 --homozyg-window-snp 65 --homozyg-window-het 1 --homozyg-window-missing 3 --homozyg-window-threshold 0.05 --homozyg-snp 65 --homozyg-kb 100 --homozyg-density 5000 --homozyg-gap 5000). The individual genome-based inbreeding coefficient, denoted as *Froh*, is defined as the fraction of total ROH length to genome effective length(Gazal et al. 2014).

### SNP densities

To check the distribution pattern of SNPs in the genomes, we adopted a method that was described by Hacquard *et al*.(Hacquard et al. 2013) Specifically, to estimate the distributions of the high- and low-SNP densities, we fitted a two-component mixture model to the observed SNP densities using the expectation-maximization (EM) algorithm (function normalmixEM, R-package mixtools). SNP densities were obtained via a sliding window of 200 kb, at steps of 2 kb, in scaffolds with lengths longer than 300kb. To identify regions with high- and low-SNP densities, a two-state hidden Markov model (HMM) was fitted on the 200-kb SNP densities using the EM algorithm, and the posterior state sequence was computed via the Viterbi algorithm (function fit, package depmixS4).

### Selective sweep identification

To detect putative selective sweeps, we searched genomic regions with higher degrees of fixation, following previously described methods(Rubin et al. 2010; Rubin et al. 2012). The numbers of major and minor allele reads observed at each variant position were counted, and SNP positions which located on non-autosomes and whose minor allele frequency was <0.05 were filtered. We then scanned the genome using sliding 100-kb windows with a step size of 50 kb. Windows with less than five SNPs were not considered. Windows with ZHp ≤–6 were retained as candidate selective sweeps.

### Salinity analyses

5 mg, 10 mg, 15 mg, 20 mg, 25 mg, 40 mg, 60 mg, 80 mg, 100 mg, 120 mg, 140 mg, 160 mg, 180 mg, and 200 mg of NaCl were weighed respectively in separate beakers. A total of 50 ml of distilled water was subsequently mixed with each quantity of NaCl to prepare saline standards. The electric conductivity (EC) value of standard saline was determined using a conductivity meter and the resultant values were used to generate the X-axis. The saline standard concentration values were used as the Y-axis. A total of 0.5 g of plant materials was weighed in a beaker and mixed with 100 ml of distilled water. After the mixture was heated using an electric stove for 30 min, the resultant solution was strained into a new volumetric flask with 25 ml of distilled water. The solution was stored in a 50-milliliter centrifuge tube and was subsequently used to determine EC values.

### Metagenomics analyses

10 fresh fecal samples from three core areas in Dafeng Natural Reserve (China) were collected immediately after defecation, snap-frozen in liquid N2, and shipped to the laboratory on dry ice. All samples were obtained from inside the feces, where there was no contact with soil. DNA was extracted from fecal samples using the Qiagen QIAamp DNA Stool Mini Kit according to the protocol for isolation of DNA for pathogen detection. DNA was eluted in a final volume of 250 μL using elution buffer and then stored at −20 °C. Sequencing and general data analyses were performed by Shanghai Majorbio Bio-pharm Biotechnology (Shanghai, China). A library was constructed with an average clone insert size of 350-bp for each sample. We compared the raw short reads with host genome data to remove the host sequence. Clean reads were subsequently obtained to assemble long contig sequences using SOAPdenovo(Li et al. 2010) during metagenomic analyses. Different Kmer frequencies were utilized to generate different assembly results, and N50 lengths were used to access the best assembly result. The metagenomes were uploaded to MG-RAST. Functional annotation of 49 metagenomes (10 from milu and 39 from published data) was performed with Hierarchical Classification using the KEGG ortholog database within MG-RAST(Meyer et al. 2008). The following parameters were used: maximum e-value cutoff of 1e-5, minimum identity cutoff of 60%, and minimum alignment length cutoff of 15 (default). The statistical analysis for KEGG function pathways were performed in STAMP(Parks et al. 2014).

